# An *ex vivo* culture model of kidney podocyte injury reveals mechanosensitive, synaptopodin-templating, sarcomere-like structures

**DOI:** 10.1101/2021.11.03.466963

**Authors:** Shumeng Jiang, Farid Alisafaei, Hong Yuan, Xiangjun Peng, Yin-Yuan Huang, Jeffrey H. Miner, Guy M. Genin, Hani Y. Suleiman

## Abstract

Chronic kidney diseases are widespread and incurable. The biophysical mechanisms underlying them are unclear, in part because material systems for reconstituting the microenvironment of the relevant kidney cells are limited. A critical question is how kidney podocytes (glomerular epithelial cells) regenerate the foot processes of the filtration apparatus following injury. Recently identified sarcomere-like structures (SLSs) with periodically spaced myosin IIA (a contractile protein) and synaptopodin (an actin-associated protein) appear in injured podocytes *in vivo*. We hypothesized that SLSs template synaptopodin in the initial stages of recovery, and tested this hypothesis by developing an *ex vivo* culture system that models both kidney physiology and pathophysiology. SLSs were observed *in vitro* for the first time as podocytes migrated out of harvested kidney glomeruli onto micropatterns of physiologically relevant proteins. SLSs emerged over two days, and cells formed foot process-like extensions from these periodically spaced proteins. SLS distributions and morphology were sensitive to actomyosin inhibitors, substrate stiffness, and extracellular matrix proteins associated with pathology. These results indicate a role for mechanobiological factors in podocyte recovery from injury, and suggest SLSs as a target for therapeutic intervention.

## Introduction

Kidney glomerular diseases result in damage to the kidney’s filtration apparatus and often cause chronic kidney disease and kidney failure with no cure ^[1]^. One barrier to devising successful treatments for these diseases is the lack of a full understanding of the biophysical mechanisms that underlie them ^[2]^. Glomerular disease may be associated with mechanobiological dysregulation of the components that comprise the three-layered filtration barrier: the endothelial cells, the glomerular basement membrane (GBM), and especially the podocytes ^[3]^. Podocytes are unique epithelial cells with hundreds of foot processes that interdigitate with those of adjacent podocytes; these are connected by slit diaphragms, unique intercellular junctions that are critical for filtration ^[4]^. However, the inability to study podocytes outside of their native microenvironment has left key gaps in our knowledge of how they maintain the intricate structures that enable filtration and of how injury and healing progress. To address this critical need, we developed a culture system that enables study of podocyte injury outside of their native microenvironment.

Previous work on podocyte mechanobiology includes a wealth of studies *in vitro* using immortalized mouse and human podocyte cell lines ^[5, 6]^, and more recently on primary mouse podocytes ^[7, 8]^, However, this work has all occurred on glass as the substrate, which is a million-fold stiffer than the ~2.5 kPa physiological range of the GBM’s elastic modulus ^[9]^. Such a mismatch is known to affect cell structure and spreading ^[10]^ and a broad range of mechanobiological responses ^[11]^. To solve these problems, a range of biomimetic platforms has thus been proposed, include culturing immortalized podocytes on soft gelatin-based ^[12]^ or Polydimethylsiloxane (PDMS) ^[10]^ substrates; curved substrates ^[13]^; or micropatterned glass substrates ^[14]^. However, these systems do not produce the structures and cell shapes observed *in vivo*, partially because they failed to sufficiently reconstitute the microenvironment that podocytes require. Due to the fact that the mechano-responsiveness of podocytes in health and disease is thus largely unknown, we sought to develop a biomimetic platform that combines primary podocytes; physiologically relevant substrate stiffnesses; physiologic or pathophysiologic ECM proteins; and physiologically relevant cell confinement.

The gap in knowledge that we sought to address relates to the mechanisms that podocytes use to repair connections to the GBM and to their adjacent podocyte neighbors. Studies of glomeruli *in vivo* have revealed that in mouse models of kidney diseases, podocytes contain the normally absent contractile protein myosin IIA in the basal aspect of the areas of foot process effacement, and the injured podocytes develop sarcomere-like structures ^[15]^. Based on these *in vivo* data, we have speculated that SLSs are associated with responses to mechanobiological cues associated with pathology, and possibly associated with podocyte migration and healing ^[8, 16]^. However, because SLSs are difficult to study *in vivo* due to their nanoscale size, and because they have not been observed in any current *in vitro* systems, this hypothesis has not been possible to test. We therefore demonstrated the usefulness of our novel *ex vivo* culture system by testing this hypothesis and identifying mechanobiological factors that collectively regulate SLSs.

## Results

### Micropatterned substrates with defined protein patterns representative of healthy and pathologic GBM

Using a freshly fabricated polydimethylsiloxane (PDMS) mold, we printed patterns of ECM proteins representative of the GBM in health and injury. To represent physiologic GBM, we used a polyacrylamide (PAAm) hydrogel of defined stiffness to microprint human laminin α5β2γ1 trimers (Lam-521, Figure 1a), the dominant laminin trimer in the GBM ^[17]^. To represent injury, Lam-521 was replaced with fibronectin (Figure 1b), which represents a GBM change observed in certain glomerular diseases and podocyte injuries ^[18]^. Using immunofluorescence staining of the hydrogel with specific antibodies to the different ECM proteins, patterns were clearly visualized as 105 µm x 7 µm microprints (Figure 1b). To microprint two different ECM substrates, we coated the hydrogel with the first ECM protein and let it dry before microprinting the second ECM protein atop the first (Figure 1a). In this way, Lam-521 micropatterns were printed atop PAAm hydrogels that were coated with either collagen IV, representative of a healthy GBM, or fibronectin, representative of abnormal GBM (Figure 1c).

**Figure 1.**
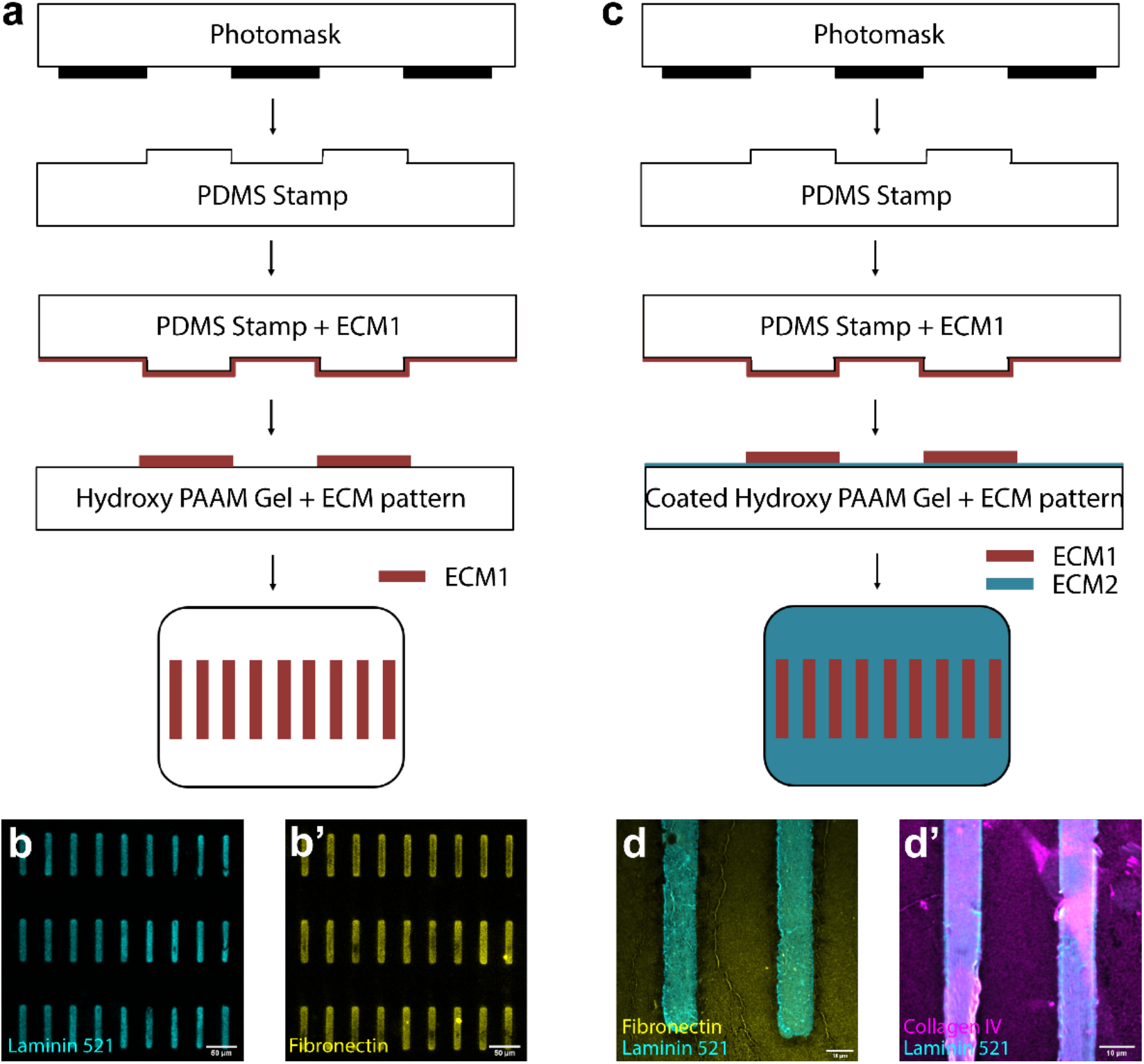
Fabrication of substrates with defined proteins patterns. a) Microprinting protocols for the fabrication of micropatterns with the desired ECM1 with the help of lithography. b) Successfully patterned laminin 521 or fibronectin on hydroxy PAAM hydrogels. c) Precoated hydroxyl PAAM hydrogel with the desired ECM2 is used instead of a plain hydrogel to enable coating both on and outside of the patterned area. d) Successfully patterned hydrogels with laminin 521 on the pattern and fibronectin or collagen IV outside of the patterned area.

### Podocytes from isolated glomeruli migrate onto micropatterned Lam-521

To test which micropatterned ECM proteins best support podocytes, we isolated mouse glomeruli and cultured them on hydrogels coated with Lam-521, collagen IV, or fibronectin. Culturing the glomeruli for 2 days followed by immunostaining using antibodies against synaptopodin, a podocyte-specific cytoskeletal marker, and human laminin α5, to identify the location of the Lam-521 micropatterns (Figure 2), revealed that the podocytes populated micropatterns coated with Lam-521 (Figure 2a) but not those coated with fibronectin (Supplementary Figure 1). Furthermore, we coated the PAAm hydrogel with Lam-521 and micropatterned protein-free patches onto the substrate using a PDMS stamp that removed Lam-521, leaving an inverse of the previous Lam-521 micropatterns interdigitated with microscale patches of exposed PAAm. Using this setup to culture glomeruli showed that podocytes migrated onto the Lam-521 coated areas, but not the bare micropatterned PAAm hydrogel areas (Figure 2b, areas outside of the red boxes).

**Figure 2.**
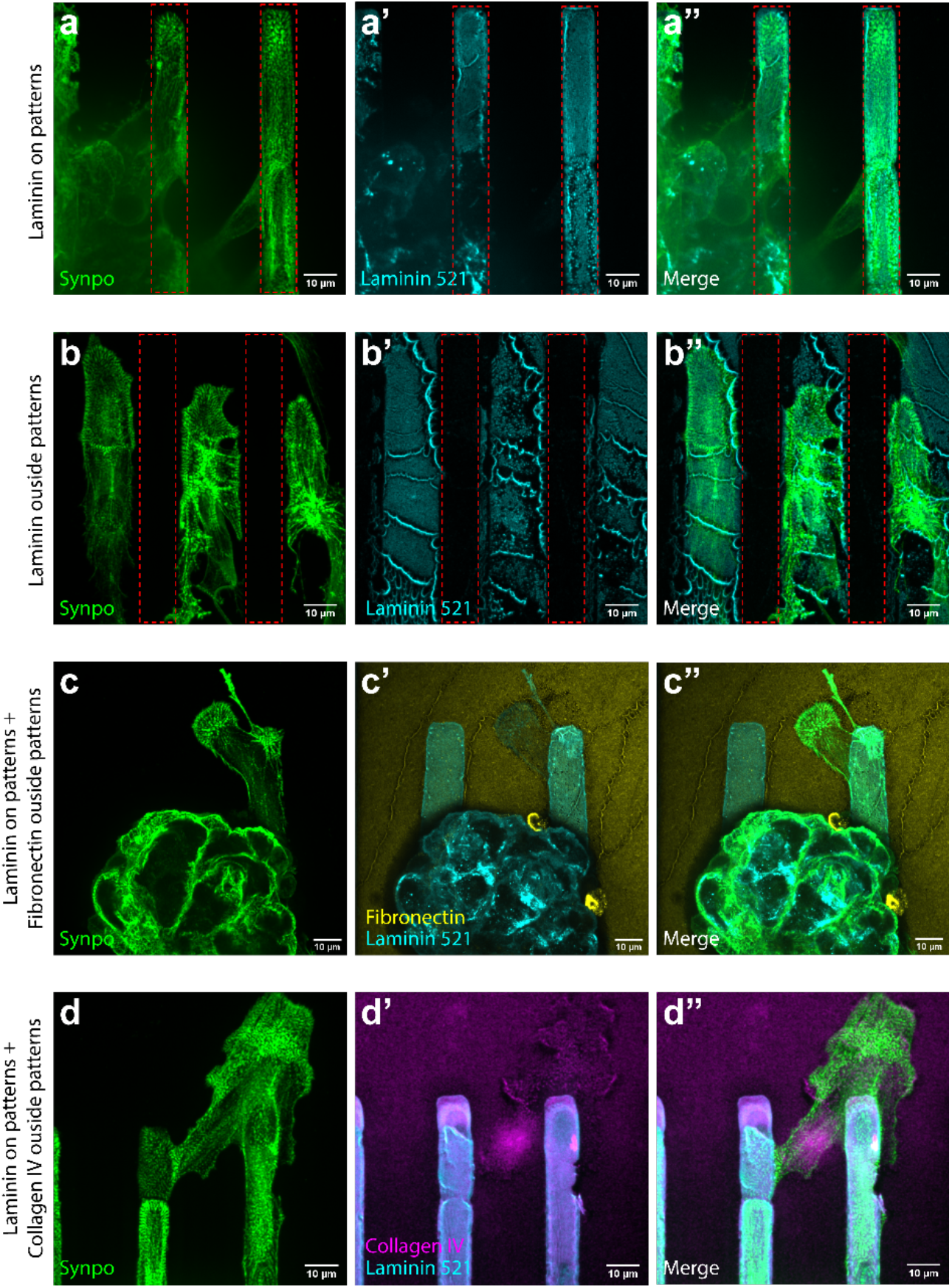
Micropatterns with combined ECM proteins reveals different stages of podocyte spreading. a) Spreading primary podocytes from an isolated mouse glomerulus can be found in the patterned area where laminin-521 is presented (marked with red boxes). b) A hydrogel precoated with Laminin-521 and printed with a blank stamp shows laminin-521 outside of the patterned area (marked with red boxes). Spreading podocytes are found only on the laminin-521. c) Podocytes spread along the micropatterned laminin-521 before sending out protrusions on the Fibronectin-enriched area outside the micropatterns. d) Primary podocytes populated the laminin-521 micropatterned area first and then spread onto the collagen IV coated area.

Next, we used PAAm hydrogels coated with fibronectin and micropatterned with Lam-521. As expected, glomeruli attached only to the Lam-521 micropatterns and podocytes only migrated onto them, whereas other cell types readily spread on both substrates (Supplementary Figure 2). Interestingly, podocytes could extend protrusions into regions covered with fibronectin (Figure 2c), but only after they migrated onto the Lam-521 micropatterns. Sometimes they migrated across regions of fibronectin to neighboring Lam-521 micropatterns (Supplementary Figure 3). Similar results were observed for collagen IV-coated PAAm hydrogels with Lam-521 micropatterns, with the majority of the podocytes confined to the Lam-521 micropatterned regions; podocytes occasionally extended processes onto collagen IV from their Lam-521 foundations (Figure 2d).

**Figure 3.**
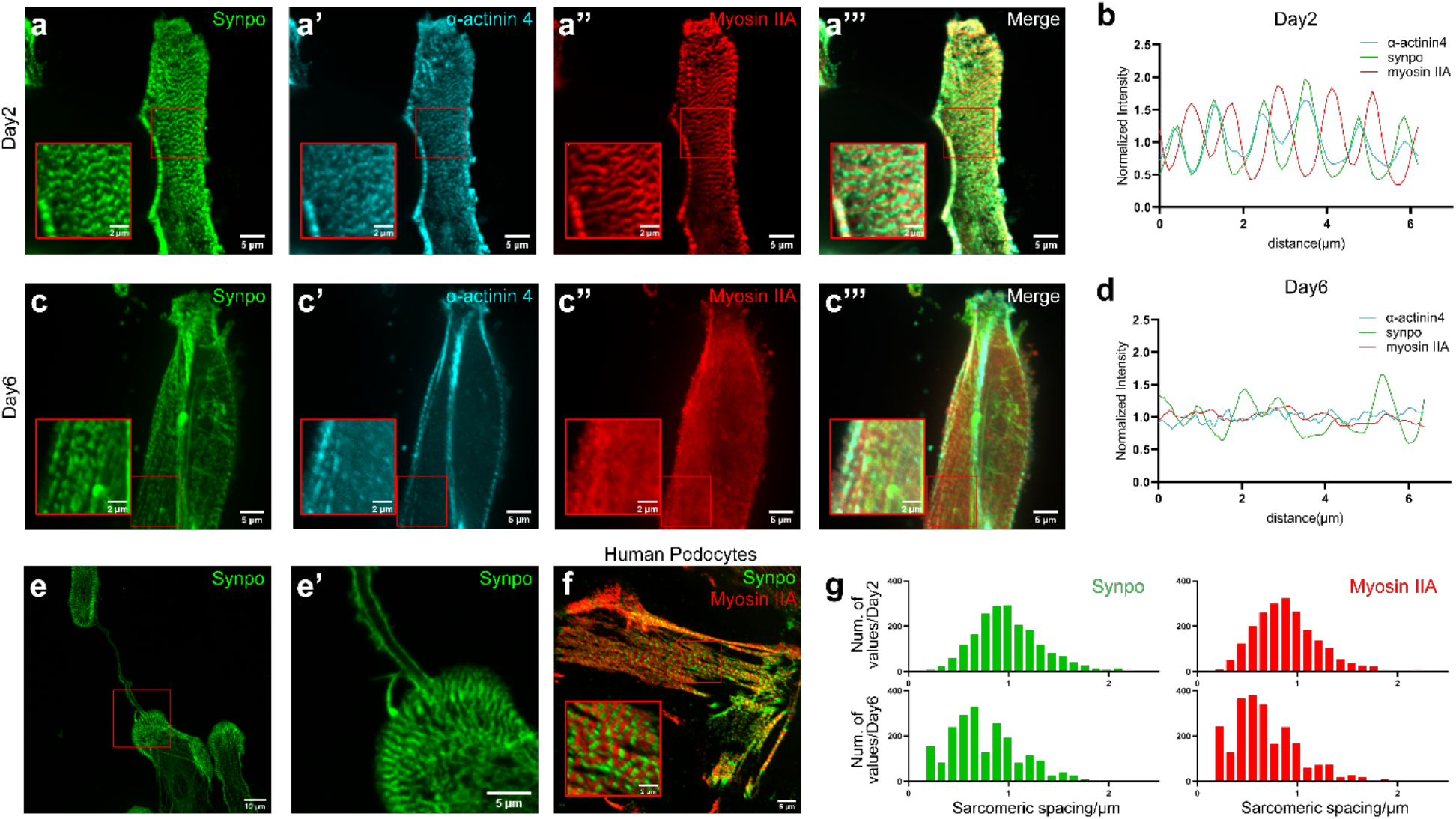
SLSs direct the early stage spreading of primary podocytes. a) SLSs could be identified after 2 days of culture, with alternating synaptopodin, α-actinin 4 and myosin IIA along the fibers. b) Fluorescence intensity mapping along the fibers’ direction shows myosin IIA alternating with co-localized synaptopodin and α-actinin 4. c) Alternating α-actinin 4 and myosin is attenuated after 6 days of culture. d) The intensity map confirms the attenuated pattern shown in c. e) Protrusions sent by the cells is positive for synaptopodin, and the width correlates to the periodic spacing of synaptopodin in the SLSs. f) SLSs could be found in human primary podocytes after 3 days of culture. g) Histogram of sarcomeric spacing represented by both synaptopodin and myosin IIA signals shows a uniform distribution of around 0.8 µm to 1 µm after 2 days of culture. The spacing was perturbed after 6 days of culture.

### SLSs are associated with healing responses in spreading podocytes and are reversible

SLSs have been described in injured podocytes *in vivo* ^[15]^, but have never been observed in cultured podocytes. Immunostaining of primary podocytes migrating out of the glomeruli onto the micropatterned hydrogels showed high numbers of SLSs two days after seeding of glomeruli onto the Lam-521 micropatterns (Figure 3). These contained two markers of contractile activity: i) the motor protein myosin IIA, and ii) synaptopodin, an actin-binding protein present in podocytes, arranged in a sarcomeric pattern of alternating synaptopodin and myosin IIA (Figure 3a). α-actinin 4, the major α-actinin isoform present in podocytes ^[19]^, was present in a striated pattern in registry with synaptopodin (Figure 3a). This periodicity was confirmed by quantifying fluorescence intensity along an axis paralleling the Lam-521 micropatterns (Figure 3b).

To assess the possibility that SLSs are representative of a healing podocyte phenotype, we studied their role in establishing connectivity to other cells. On substrates that presented cells with fibronectin, which is representative of injury, outside of the patterned Lam-521 area, podocytes with SLSs sent out protrusions over the fibronectin (Figure 2c, Supplementary Figure 3). These processes resemble those observed in development in native podocytes ^[20]^. Some processes extended to connect podocytes across fibronectin patterns that were several cell widths apart (Figure 3e). An additional aspect of the SLSs relevant to healing was an apparent role of the periodically spaced synaptopodin in guiding the formation, spacing, and width of the protrusions (Figure 3e).

We next asked whether SLSs represent a transient or a permanent change in the podocytes. To address this question, we followed podocytes over a prolonged culture interval of 6 days. When compared to the 2-day cultures, the SLSs in 6-day cultures showed less pronounced synaptopodin (Figure 3b, Supplementary Figure 4), as evident from the spatial intensity map (Figure 3d) and from the histogram of sarcomeric spacing (Figure 3g). This suggests that synaptopodin in SLSs became attenuated over time.

**Figure 4.**
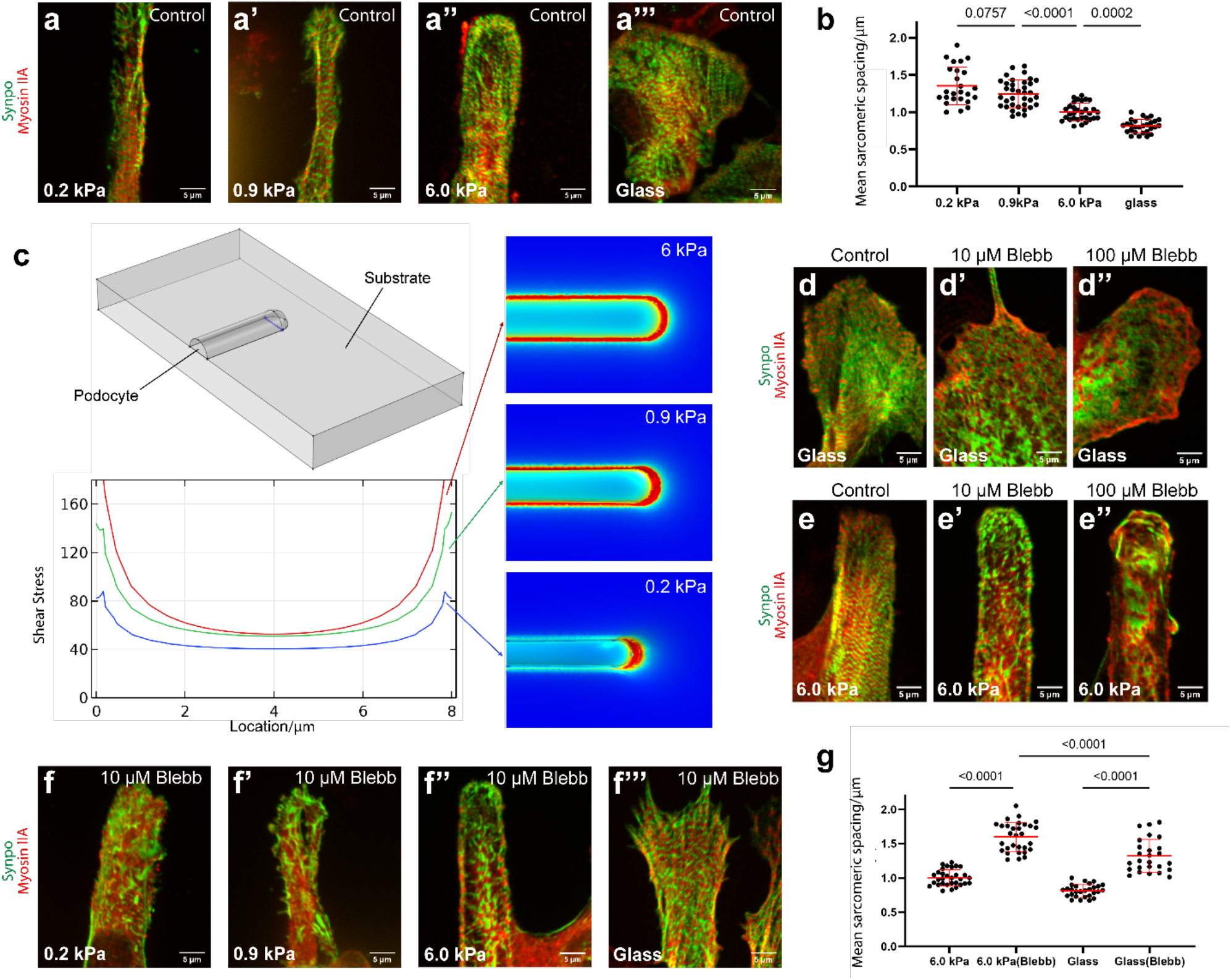
The SLSs of podocytes are dependent upon contractility and are sensitive to stiffness. a, b) The uniformly distributed 1µm-long SLSs elongated to 1.4µm when the substrate stiffness was decreased from 6.0 kPa to 0.2 kPa. In contrast, the extremely stiff glass substrate shortened this spacing to 0.8µm. c) Modeling of cell contractility on different substrates shows increased contractility on stiffer hydrogels. d, e) Myosin inhibition by blebbistatin caused dramatic loss of SLSs inside podocytes cultured either on glass (d) or on patterned hydrogels (e). f) Combining the effect of myosin inhibition with that of a softer substrate caused the loss of SLSs with a lower concentration of blebbistatin. g) For podocytes on the stiffest substrate, myosin inhibition led to wider sarcomeric spacing.

To confirm that SLSs are relevant to human podocytes, we applied our protocols to freshly harvested glomeruli taken from human kidney nephrectomy samples and seeded them onto the Lam-521 micropatterned hydrogels. Similar to the mouse glomeruli, human glomeruli attached to the Lam-521 micropatterns, although they took longer to attach when compared with mouse glomeruli. After 3 days of culture, sarcomeric structures could be observed in podocytes migrating from human glomeruli (Figure 3f), confirming that SLSs are present in primary human podocytes when migrating onto the micropatterns.

### SLSs are mechanosensitive

We next asked whether SLSs are sensitive to their mechanical microenvironment. PAAm hydrogels with elastic moduli in the physiologic range (0.2 and 0.9 kPa) and in pathophysiologic ranges (6.0 kPa, representative of GBM stiffening under hypertension or glycation ^[21]^, and cover glass, as a fully rigid substratum) were micropatterned or coated with Lam-521. After 2-day culture of glomeruli, SLSs were evident in podocytes on substrata of all moduli, but the spacing of the sarcomeric patterning was wider on the softer hydrogels than on the stiffer hydrogels and cover glass (Figure 4a). The periodic synaptopodin distribution from 120 cells over three replicate experiments showed substantial differences between softer hydrogels (0.2 kPa or 0.9 kPa) and stiffer hydrogels or glass (Figure 4b).

### SLSs require actomyosin contractility

Given that mechanosensitivity and mechanosensing in other cell types arise from actomyosin contractility ^[22, 23]^, we next asked whether SLSs can be controlled by modulating myosin II using the drug blebbistatin. In cells such as smooth muscle cells and fibroblasts, reductions in substrate stiffness reduce the resistance to actomyosin contractility and can thereby destabilize stress fibers and focal adhesions that require force for stability ^[24]^.

Our numerical simulations revealed that stresses were highest along the periphery of cells and at their distal ends (Figure 4c). Stresses also increased with substrate modulus (Figure 4c) and were attenuated by reduced cell contractility (Supplementary Figure 5). Following a standard model of mechanosensitive stabilization and growth of stress fibers ^[25]^, these simulations thus predicted that stress fibers disassemble in the interior of cells and at the proximal ends when actomyosin contractility is attenuated either by actomyosin inhibition or by reducing substrate modulus. Simulations also predicted that cell width decreases with decreasing substrate modulus (Figure 4c).

To test these predictions, we first applied two concentrations of blebbistatin to podocytes cultured on substrates with high stiffness (6.0 kPa PAAm hydrogels and cover glass), micropatterned or coated with Lam-521. Myosin II inhibition via blebbistatin disrupted the sarcomeric structures of SLSs in a way that depended upon dose, substrate stiffness, and position (Figure 4d-e), consistent with the model predictions. On both substrates, 10 µM blebbistatin caused aggregation of synaptopodin in the center of the cells and increased sarcomeric spacing in SLSs that persisted at the cell periphery (Figure 4d-e), as predicted by the model. Higher doses (100 µM) of blebbistatin nearly eliminated SLSs (Figure 4d-e). Further experiments applied a milder myosin inhibition using 2 µM blebbistatin to verify this shear lag effect. This was insufficient to disrupt the SLSs at the cell periphery but sufficient to disrupt those in the centers of the cells, and the corresponding distribution of SLSs in podocytes followed the model’s predictions (Supplementary Figure 6).

A second prediction of our model was that soft substrates lead to lower cell contractility and thus enhance the effects of actomyosin inhibition. To test this, we explored whether a minimum stiffness threshold was required for formation of SLSs in cells treated with 10 nM blebbistatin. In podocytes treated with 10 nM blebbistatin and cultured on 6.0 kPa hydrogels or glass substrates, SLSs formed, but with wider sarcomeric spacing (Figure 4f-g). When this treatment was applied to podocytes cultured on 0.2 kPa or 0.9 kPa hydrogels, representative of under-stressed GBMs in hypotensive conditions due to the nonlinearity of GBM ^[26]^, SLSs failed to form, with no sarcomeric structure evident (Figure 4f). This supported the hypothesis that actomyosin-based mechanosensation governs formation of SLSs, and that a threshold of developed tension is required.

Finally, our model predicted that cells would contract laterally on substrates of lower stiffness: the reduced contractile stresses associated with low substrate stiffness would be outweighed by the decreased resistance to lateral contraction. This prediction was borne out by our experiments, with cells showing narrower bodies on substrates of lower modulus (Figure 4a).

### SLS formation can be rescued through removal of actomyosin inhibition

To test the hypothesis that mechanobiological factors alone can drive SLS formation, we asked whether SLSs susceptible to actomyosin inhibition are reversable after washing out blebbistatin. After 2 hours of inhibition, cells were left to recover for another 2h or 1 day, then stained for synaptopodin and myosin IIA. SLSs appeared as early as 2 hours following removal of blebbistatin, although some condensates of synaptopodin were still evident. After 1 day of recovery, these condensates disappeared and SLSs returned (Supplementary Figure 7).

## Discussion

Our micropatterned culture system enabled the visualization of SLSs, which had previously been observed only in injured podocytes *in vivo*. Results demonstrated SLSs in primary podocytes migrated from both mouse and human glomeruli, and establish SLSs as mechanosensitive.

Our results further suggest that SLSs are associated with a healing phenotype, with SLSs linked to conditions associated with podocyte healing or anti-detachment responses. Bands rich in myosin IIA were found to alternate in SLSs with bands rich in both synaptopodin and α-actinin 4. A role in healing is further suggested, because dominant mutations of α-actinin 4 are associated with diminished podocyte injury resistance, including focal segmental glomerulosclerosis ^[27]^. One possibility is that SLSs template periodic synaptopodin for podocyte healing and recovery, as suggested by the synaptopodin-rich processes we observed; these are reminiscent of the foot processes that are required for normal podocyte-podocyte connectivity and podocyte function. Although no specific pathogenic phenotype has been observed in synaptopodin knockout mice ^[8]^ at baseline, the loss of synaptopodin exacerbates both drug-induced and genetic kidney injuries ^[8, 16]^. Our *ex vivo* system may be useful for further clarifying synaptopodin’s roles in podocyte injury and kidney glomerular disease.

Our observation that SLS band spacing increased with increasing substrate stiffness increases suggests that they are, similar to SLSs in other cells, mechanosensitive. This is consistent with observations of mechanosensitive formation of stress fibers observed in immortalized podocytes on soft gelatin substrates ^[12]^. The formation and organization of SLSs is also interesting: To achieve the ideal spacing in the SLSs for the templating of synaptopodin, there may be different mechanisms compared to the organization of sarcomeres in the myofibrils of muscle cells, especially considering the fact that sarcomeric spacing is usually between 1.5 and 3.5 µm in myofibrils, whereas we determined the sarcomeric spacing in SLSs to be between 0.8 and 1 µm in spreading podocytes during culturing on hydrogels with physiologic stiffness.

In addition to templating, SLSs may serve as a buffer against mechanical perturbations that could cause detachment from the GBM and loss of podocytes. The sarcomeric patterns of SLSs are reminiscent of sarcomeric contractile units in muscle ^[28]^ and contractile non-muscle cells ^[29]^, including contractile actin cables that comprise the ventral stress fibers and the transverse arcs ^[30]^. Like stress fibers in fibroblasts ^[29]^, SLSs in primary podocytes aligned in the direction of maximum stress, at a density that correlated with the amplitude of stress, and may serve to stabilize motor clutch bonds to the ECM ^[23]^. SLS alignment with the direction of cell spreading might power cell spreading and colonization of the GBM surrounding the glomerular capillaries.

The micropatterns used in this study were rectangles of approximately 20 µm in length, with a length:width ratio of 14:1 to 8:1. This size allowed the spreading of podocytes on the 2D hydrogel surfaces, while part of the podocytes remained in contact with the glomerulus. Thus, podocytes remained connected to their native 3D structure while extending into confined patterns that mimicked the constraints of the GBM. One limitation of our system is that the micropatterns limit the ability of podocytes to form foot processes and slit diaphragms, and future work should address this inadequacy. Extensions of this technology via nanopatterning may better replicate the cell microenvironment and perhaps spur the formation of foot processes. The GBM has a curvature that, although large in radius compared to the size of a podocyte foot process, may nevertheless be a factor in podocyte mechanobiology. Incorporating curvature with microprinting via surface engraving technologies may enable exploration of these factors. Finally, fluid shear stresses are an additional factor that is likely important in podocyte homeostasis, and the addition of microfluidics to the system may constitute an important step forward. However, even with these limitations, our culture system has served to identify and validate essential biophysical mechanisms underlying SLSs, including mechanobiological factors that serve as potential therapeutic targets. SLSs may be a transient feature of podocyte healing or an advantageous response to injury and thus a target for intervention in kidney injury and disease.

## Materials and Methods

### Activation of cover glasses

To enable firm attachment of PAAm hydrogels to cover glasses, the cover glasses were washed twice in NaOH (0.1M) for 5 min, rinsed with ddH2O, and dried before applying a thin layer of 3-(trimethoxysilyl)propyl acrylate for 1 hour at room temperature. They were then washed again in ddH2O and dried under nitrogen flow ^[31]^.

### Preparation of hydroxy-PAAm hydrogels with defined elastic modulus

was achieved by crosslinking precursors on activated cover glasses via modification of established protocols ^[32]^. Hydroxy-PAAm hydrogels were prepared using acrylamide (3.2-6.4% w/w in HEPES, pH 7.4), bis-acrylamide (0.03-0.16% w/w in HEPES, pH 7.4) and N-hydroxyethyl acrylamide (HEA) (1.3% w/w in HEPES, pH 7.4) together with ammonium persulfate (APS) and N-tetramethylenediamine (TEMED). After incubating the pre-crosslinking solution (i.e., the acrylamide, the bis-acrylamide and the N-hydroxyethyl acrylamide) for 30 minutes under vacuum to remove oxygen and thus prevent oxidation, the crosslinking agents were added and incubated for another 30 minutes at room temperature. For 5 mL of pre-crosslinking solution, 2.5 µl of TEMED and 25 µl of 10% APS were added. To shape the hydrogel as a thin layer and protect it from oxidation, a clean, non-activated cover glass was placed on top. After the hydrogel solidified, the sample was washed 3x with ddH2O and left at 4°C. To control the stiffness of the hydrogel, both acrylamide and bis-acrylamide final concentrations were altered, with acrylamide varied from 3.2% to 6,4% w/w, and bis-acrylamide varied from 0.03% to 0.16% w/w. The moduli of hydrogels were measured using atomic force microscopy ^[33]^, and were controlled to range in stiffness from 0.9 - 6.0 kPa, a range that encompasses that reported for GBM ^[34]^.

### Micropattern stamps for microprinting

A silicon master with the desired micropattern design was prepared using standard lithographic techniques ^[35]^. A PDMS stamp was prepared by mixing PDMS and a crosslinking agent at a 10:1 ratio. PDMS was degassed under vacuum to remove air bubbles. The molding mixture was then added to a container in the presence of the silicon master template, set inside an oven (VWR) at 60°C for 2 hours to initiate the crosslinking. Next, the PDMS stamp was cleaned by sonication (Branson) for 20 min, washed in 50% ethanol solution before being drying under nitrogen flow followed by plasma cleaning for 2 min (Harrick Plasma). Finally, the PDMS stamp micropattern areas were soaked in different ECM solutions (50 µg/ml Laminin 521, Fibronectin, or Collagen IV in PBS) and left to set for 1 hour at room temperature.

After air-drying the ECM proteins atop the PDMS stamp, excess solution was removed and the stamp was dried further using nitrogen flow. The freshly prepared hydroxyl-PAAm hydrogel was also dried out by treatment with nitrogen flow. To microprint one ECM: i) the PDMS stamp was turned so that the patterned-surface faced the hydroxyl-PAAm hydrogel surface, ii) the stamp was gently placed onto the center of the hydroxyl-PAAm hydrogel and left there for one hour at room temperature, and finally iii) the PDMS stamp was removed carefully and the hydrogel was washed 3x with PBS to remove unbound proteins.

For micropatterns with combined ECM proteins, the protocol was the same except that ECM solutions were added on top of the hydroxyl-PAAm hydrogel and left for one hour at room temperature before drying and the subsequent application of the above microprinting protocol.

### Isolation of mouse glomeruli

Mouse glomeruli were isolated using an established differential adhesion method ^[36]^ with modifications. Briefly, kidneys were collected, minced, and digested in collagenase A solution in HBSS (Gibco 24020-117, 1mg/ml) at 37°C for 15 min. The collagenase A digestion was stopped by adding an equal volume of DMEM with 10% FBS. Next, the suspension containing the dissociated kidney fragments was passed through three differently sized cell strainers (100-µm, 70-µm then 40-µm, MIDSCI) and the glomeruli-enriched tissue fragments were collected on top of the 40-µm cell strainer. These were then placed onto 10 cm tissue culture dishes (TPP) for 1-2 min to allow the tubular segments to adhere before collecting the glomeruli left in suspension. For higher glomerular purity, the adhesion step was repeated twice. Finally, the suspension was spun down and the glomeruli were re-suspended in primary podocyte culture medium and directly used for the downstream applications.

The mouse podocyte culture medium was prepared as reported earlier ^[37]^. Briefly, for 648 mL of podocyte culture medium, we mixed: i) 3T3L1 conditional media (300 mL), which was the supernatant arising from culture of the 3T3L1 cell line for 3 days at 37°C in DMEM culture medium (Gibco 11965-084) containing 10% fetal bovine serum (Sigma F7524) and 1% penicillin-streptomycin (Sigma P4333); ii) low glucose DMEM (204 mL); iii) Ham’s F-12; iv) L-Glutamine (102 mL, Lonza); v) fetal bovine serum (30 mL); vi) penicillin-streptomycin (6 mL) (Sigma P4333) and vii) insulin-transferrin-selenium liquid media supplement (6 mL) (Invitrogen 41400045).

### Isolation of human glomeruli

Kidney nephrectomy tissue was minced, passed through a 250 µm metal cell strainer (MIDSCI) using 1x HBSS solution (Gibco 24020-117), and collected on a 70-µm cell strainer. The glomeruli-rich suspension was spun down at 2000g (g = 9.81 m/s^2^) for 5 min and rinsed by passing it through the 70-µm strainer using HBSS. Finally, the glomeruli on top of the 70-µm cell strainer were rinsed and re-suspended in human primary podocyte culture medium, consisting of DMEM/F12 medium, (Thermofisher 11320-033) with 1% v/v insulin-transferrin-selenium (Invitrogen 41400045), 20% v/v fetal bovine serum (Sigma F7524) and 1% v/v penicillin-streptomycin (Sigma P4333). These were directly used for the downstream applications ^[38]^.

### Glomerular culture and podocyte spreading on micropatterned hydrogels

Before culturing the glomeruli, micropatterned hydrogels were moved inside a 6-well culture plate and washed thoroughly with HBSS. After the isolation, glomeruli suspended in podocyte culture medium were placed on the micropatterned hydrogel. For a single hydrogel, ~100 µl of the glomerular suspension was added and cultured at 37°C, 5% CO_2_ and 95% humidity. Additional podocyte culture medium (2 ml) could be added to the culture after 10 hours of attachment.

### Immunofluorescence staining

After spreading, cultured podocytes on the hydrogels or glass were fixed in 4% PFA (Electron Microscopy Sciences 15712) for 10 minutes at room temperature followed by washing with 1x PBS, three times for 6 minutes each, then permeabilized using 0.05% Triton X-100 for 10 minutes at room temperature. Next, samples were blocked using 2% bovine serum albumin (BSA, Sigma-Aldrich A7906) for 30 min at room temperature before applying the primary antibodies overnight at 4°C. Next, samples were washed with 1x PBS, three times for 6 minutes each before incubating them with the secondary antibodies in PBS for 1 hour at room temperature. Finally, the samples were washed in 1x PBS, three times for 6 minutes and prepared for mounting using Invitrogen Slow Fade mounting medium (Invitrogen S36917). The cover glass holding the sample (hydrogel plus cells) was bonded to a second cover glass using nail polish.

The primary antibodies used were myosin IIA (Biolegend, 909801, rabbit anti–mouse), myosin IIA (Abnova, clone 3C7, mouse anti–human, NH2-terminus), synaptopodin (Synaptic Systems, 163004, guinea pig anti-mouse), Laminin α5 (clone 4C7, mouse anti–human, Engvall et al., JCB 1986), Fibronectin (Sigma-Aldrich, F3648, Rabbit anti–human), Collagen IV (SouthernBiotech, 1340-01, Goat anti–human). Alexa Fluor 555 Phalloidin (Thermo Fisher Scientific, A34055) was used for actin staining. For mouse myosin antibodies (clone 3C7), samples were antigen retrieved in TE buffer (pH 9.0) at 65°C for 4 hours before the permeabilization step.

### Confocal microscopy

Imaging was performed on a Zeiss LSM 880 confocal microscope (equipped with a unique scan head incorporating a high-resolution galvo scanner along with two PMTs and a 32-element spectral detector as well as a transmitted light PMT for DIC imaging) or a Nikon spinning disk confocal microscope (equipped with Yokagawa CSU-X1 variable speed Nipkow spinning disk scan head, Andor Zyla sCMOS cameras and an LED-based DMD system for ultrafast photo-stimulation). Images were taken using a 10X objective for micropatterns with a single protein and using a 60X or 100X oil immersive objective for other images.

### Myosin II inhibition with blebbistatin

Blebbistatin (EMD Milipore 203389-5) was used to inhibit the function of myosin II in the podocytes. At the second day of primary podocyte culture, blebbistatin (50 µM stock solution) was added directly into the podocyte culture medium for a final concentration ranging from 2 nM to 100 nM. After 2 hours of treatment, the samples were washed with PBS and immediately fixed and stained as mentioned above.

### Numerical simulations

A finite element model of a cell process contracting isotropically atop an elastic substrate was studied. The cell and substrate were approximated as isotropic continua. The cell had an elastic modulus of 1.0 kPa and a Poisson ratio of 0.3. The substrate had an elastic modulus that was varied from 0.2 to 6.0 kPa and a Poisson ratio of 0.3. The cell process was given the dimensions shown in Figure 4c. The substrate base was fixed, and the sides were traction free. Convergence was achieved when the substrate was discretized with 67354 quadratic elements, and the cell with 50573 quadratic elements. The cell’s reference configuration contracted by 10% in x direction, while 8% in y direction and 2% in z direction, considering the main contractility should follow the direction of the stress fiber which is in the x direction according to Figure 3a, but this contraction was resisted by elastic stresses stored in the cell and substrate. Simulations were performed using Comsol (Comsol, Inc., Burlington, MA).

### Quantitative microscopy of sarcomeres

Sarcomeric structures were measured using ImageJ (v1.53e with Bio-Formats plugin). Briefly, fluorescence intensity maps were acquired by plotting the intensity along a line drawn perpendicular to the sarcomere structure. For histogram, spacing of each sarcomere is calculated by recording the peak location in the intensity map, and over 2000 SLSs is recorded for each group. For averaged sarcomeric spacing, the mean values were calculated for each intensity map (i.e., each stress fiber), while at least 25 stress fibers were included for each group.

## Acknowledgments

This work was funded in part by the Office of the Vice Chancellor for Research at Washington University in St. Louis, by a Pilot and Feasibility grant from the Diabetes Research Center (NIH P30DK020579) to HS, by NIH grants (R01DK058366 and R01DK078314) to JHM, and by the National Science Foundation Center for Engineering Mechanobiology (CMMI 1548571) to GMG.

## Conflict of Interest

There is no conflict of interest.

## Notes

### Competing Interest Statement

The authors have declared no competing interest.

