## Supplemental data for "An *ex vivo* culture model of kidney podocyte injury reveals mechanosensitive, synaptopodin-templating, sarcomere-like structures"

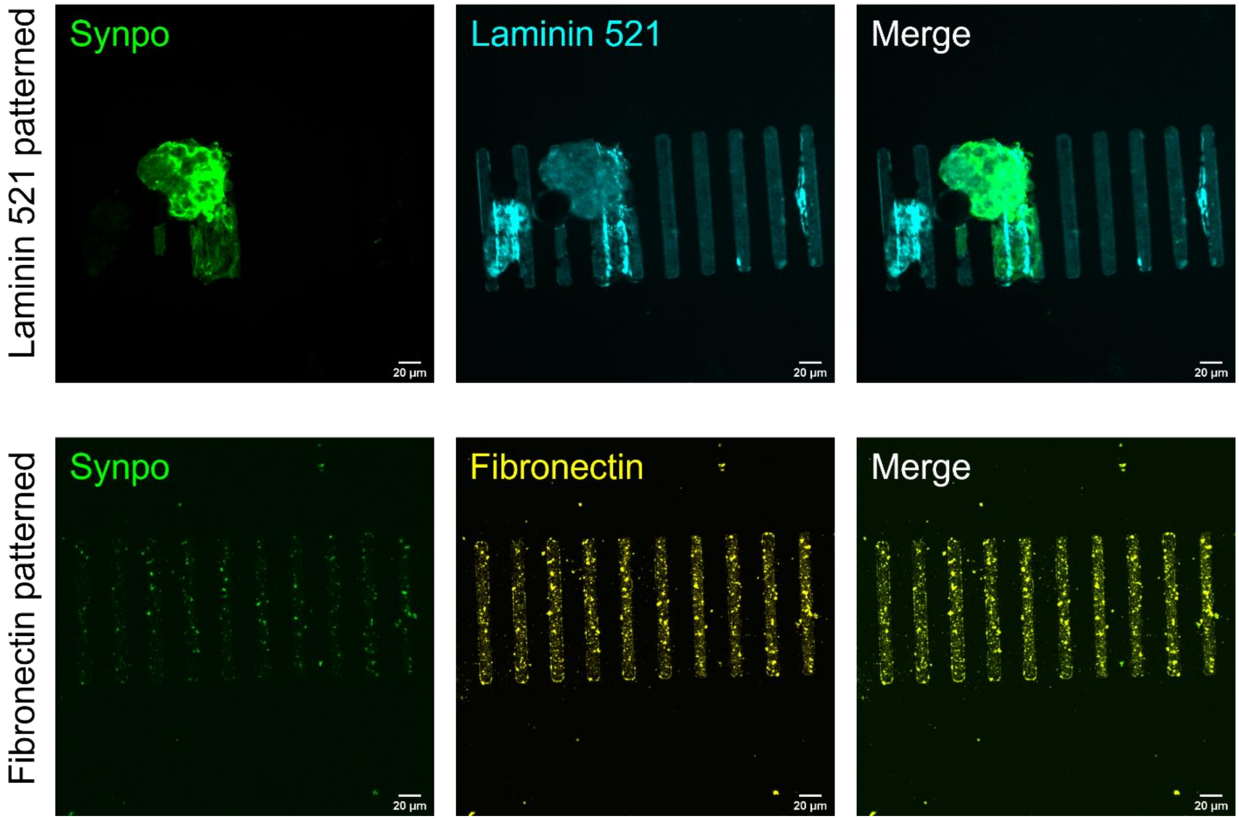


**Supplementary Figure 1** Unlike Laminin 521 micropatterns, Fibronectin micropatterns are not efficient at enabling glomerulus/podocyte attachment on the hydrogels.

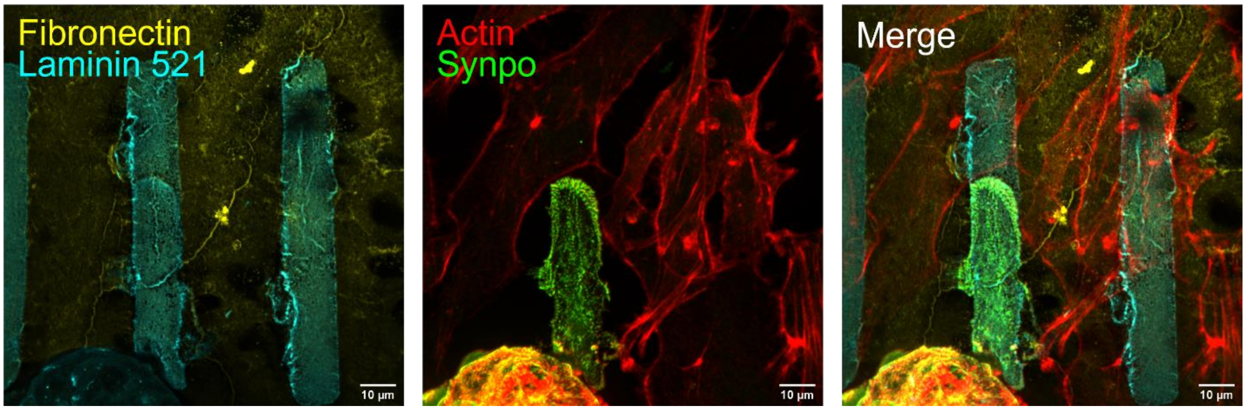


**Supplementary Figure 2** The preferential adhesion/migration on laminin-521 is specific for podocytes, as there are many cells without synaptopodin attached to the fibronectin between the laminin-521 micropatterns.

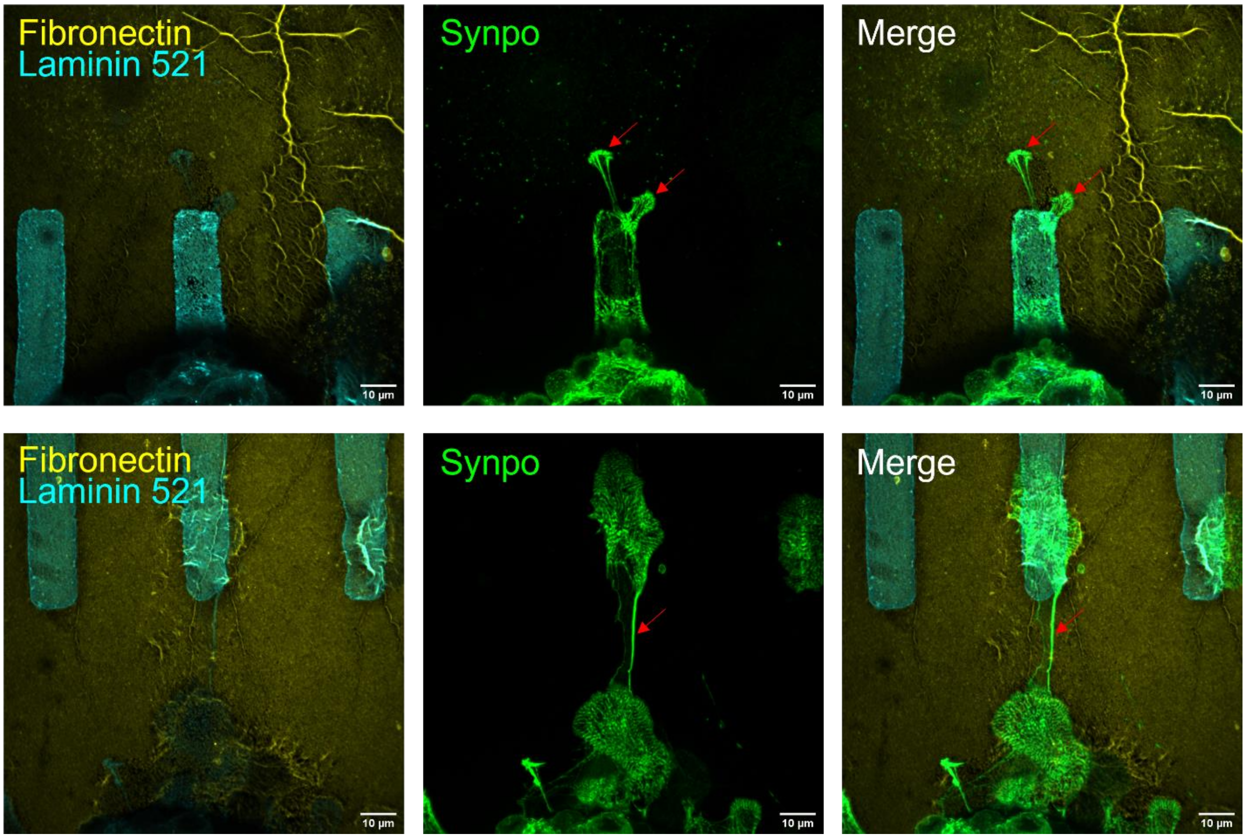


**Supplementary Figure 3** When cultured on Laminin 521 micropatterned hydrogels with Fibronectin outside the micropatterns, the podocytes send out synaptopodin-positive protrusions (Marked by red arrows) which seems to enable migration to another nearby micropattern.

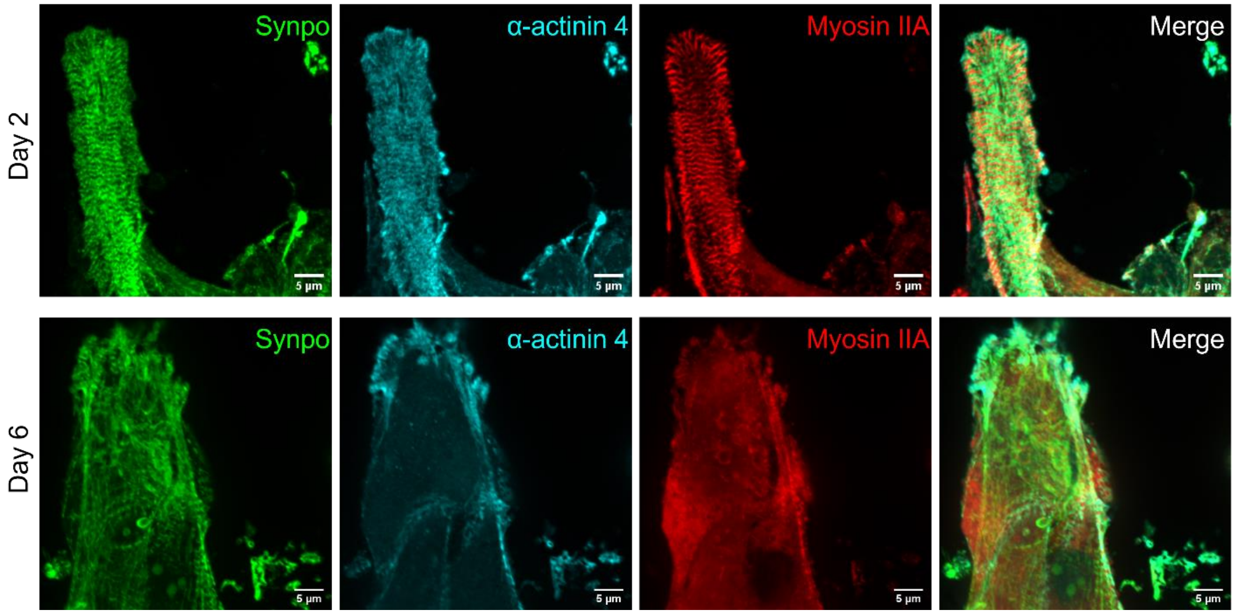


**Supplementary Figure 4** Additional images show podocytes with SLSs on day 2, but most of them are lost by day 6.

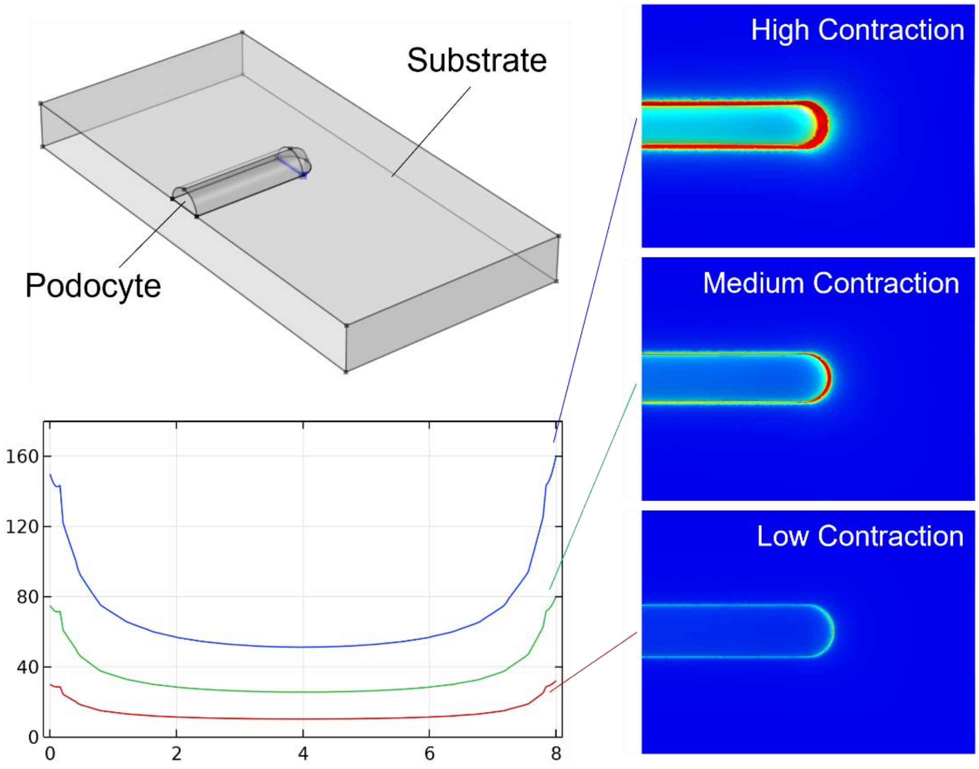


**Supplementary Figure 5** Modeling the shear stress at the adhesion surface shows dramatically reduced shear stress when the contractility inside of the cells is inhibited.

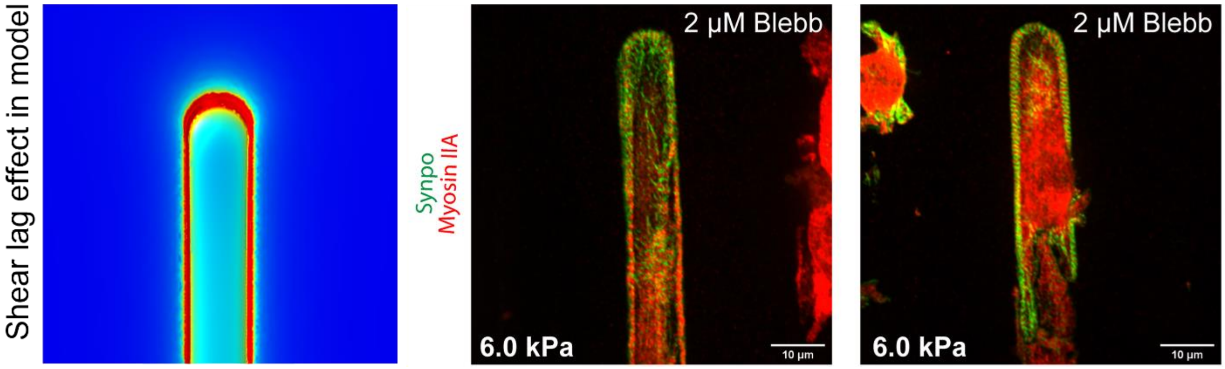


**Supplementary Figure 6** Mild myosin inhibition is not enough to eliminate all the SLSs inside of the spreading podocytes, but SLSs in the center of the cells were disturbed due to shear lag effects.

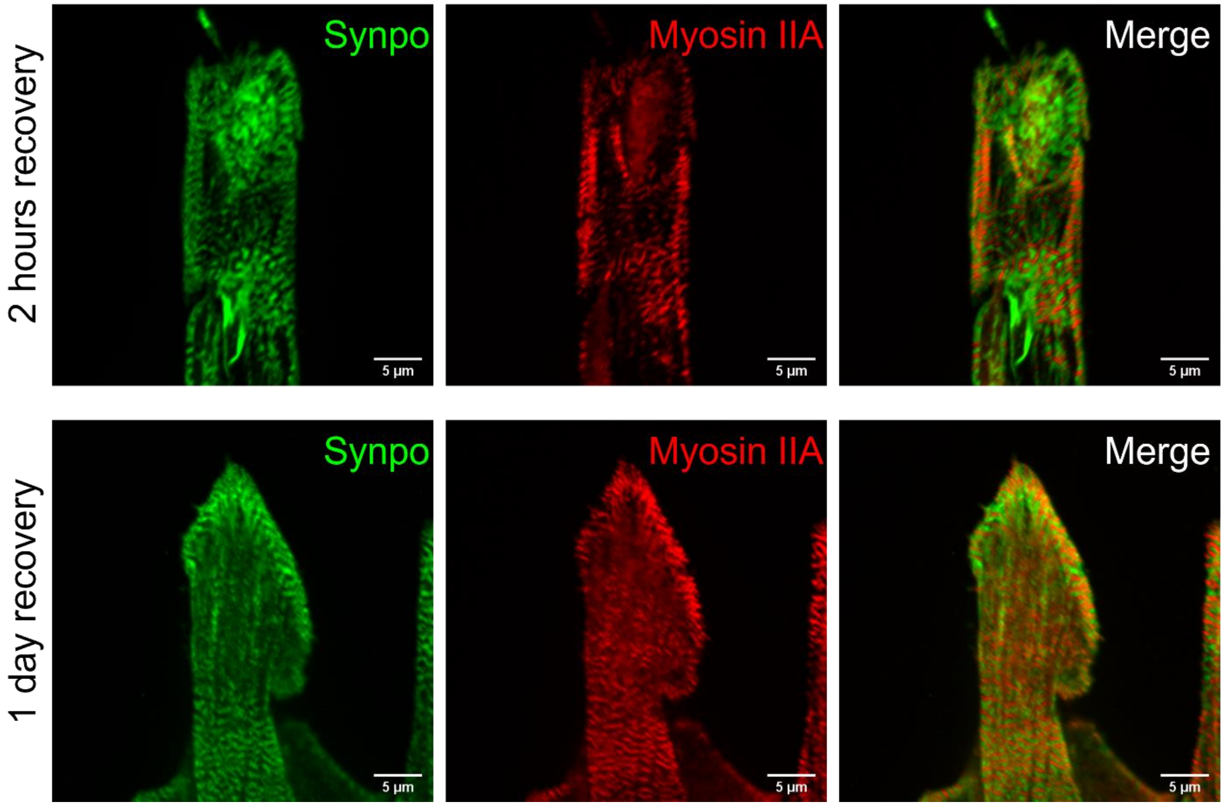


**Supplementary Figure 7** Blebbistatin wash-out lead to the regeneration of SLSs in podocytes within 2 hours. Longer (1 day) recovery allowed restoration of the SLSs even in the center of the cells.
